## Supplementary material for "Vaccinia virus infection induces concurrent alterations in host chromatin architecture, accessibility, and gene expression"

| Biological Replicate | Sample | Total read-pairs | Hi-C contacts | Contacts (Per condition) | Loops called |
|---|---|---|---|---|---|
| 1 | 12hr Control | 559,483,569 | 256430862 | 783,687,719 | 10146 |
| 2 | 12hr Control | 886,507,172 | 534469766 | | 17667 |
| 1 | 12hr Infected | 777,613,496 | 270746905 | 712,124,938 | 8771 |
| 2 | 12hr Infected | 770,418,243 | 463757732 | | 15975 |
| 1 | 18hr Control | 761,567,093 | 327008067 | 702,267,757 | 13569 |
| 2 | 18hr Control | 610,190,915 | 381729147 | | 17416 |
| 1 | 18hr Infected | 646,594,007 | 279263882 | 653,725,550 | 10064 |
| 2 | 18hr Infected | 655,739,644 | 408610556 | | 15379 |
| 1 | 24hr Control | 725,697,285 | 449311017 | 846,643,183 | 17871 |
| 2 | 24hr Control | 619,039,813 | 405023378 | | 17092 |
| 1 | 24hr Infected | 781,644,457 | 462522234 | 799,501,451 | 14718 |
| 2 | 24hr Infected | 607,026,371 | 387415043 | | 13988 |

*Table S 1: Hi-C data summary.*

| Biological Replicate | Sample | Total read pairs | ATAC-seq peaks in assembled chromosomes | FRiP score |
|---|---|---|---|---|
| 1 | 12hr Control | 42,598,767 | 126,980 | 0.69 |
| 2 | 12hr Control | 22,250,266 | 148,912 | 0.51 |
| 1 | 12hr Infected | 39,289,957 | 98,872 | 0.47 |
| 2 | 12hr Infected | 65,728,373 | 126,769 | 0.54 |
| 1 | 18hr Control | 17,094,491 | 114,885 | 0.75 |
| 2 | 18hr Control | 64,577,918 | 135,385 | 0.61 |
| 1 | 18hr Infected | 29,811,600 | 57,763 | 0.46 |
| 2 | 18hr Infected | 58,849,770 | 75,849 | 0.38 |
| 1 | 24hr Control | 21,883,403 | 110,433 | 0.72 |
| 2 | 24hr Control | 78,668,288 | 162,898 | 0.59 |
| 1 | 24hr Infected | 35,202,941 | 835,69 | 0.38 |
| 2 | 24hr Infected | 86,633,067 | 140,331 | 0.43 |

*Table S 2: ATAC-seq data summary*

| Biological Replicate | Sample | Total read pairs | Total aligned read pairs | Genes with 5X or greater read depth |
|---|---|---|---|---|
| 1 | 12hr Control | 16,724,888 | 14,188,215 | 14264 |
| 2 | 12hr Control | 7,853,774 | 6,475,281 | 12081 |
| 1 | 12hr Infected | 40,441,286 | 34,712,486 | 17491 |
| 2 | 12hr Infected | 14,313,472 | 11,975,560 | 15458 |
| 1 | 18hr Control | 14,881,111 | 12,319,322 | 15178 |
| 2 | 18hr Control | 13,288,297 | 11,465,064 | 15179 |
| 1 | 18hr Infected | 9,962,273 | 8,520,981 | 14161 |
| 2 | 18hr Infected | 17,880,406 | 15,165,670 | 15599 |
| 1 | 24hr Control | 19,478,850 | 16,443,010 | 15774 |
| 2 | 24hr Control | 19,602,211 | 16,540,109 | 15879 |
| 1 | 24hr Infected | 17,689,469 | 14,446,357 | 15318 |
| 2 | 24hr Infected | 17,537,009 | 14,926,234 | 15283 |

*Table S 3: RNA-seq data summary*

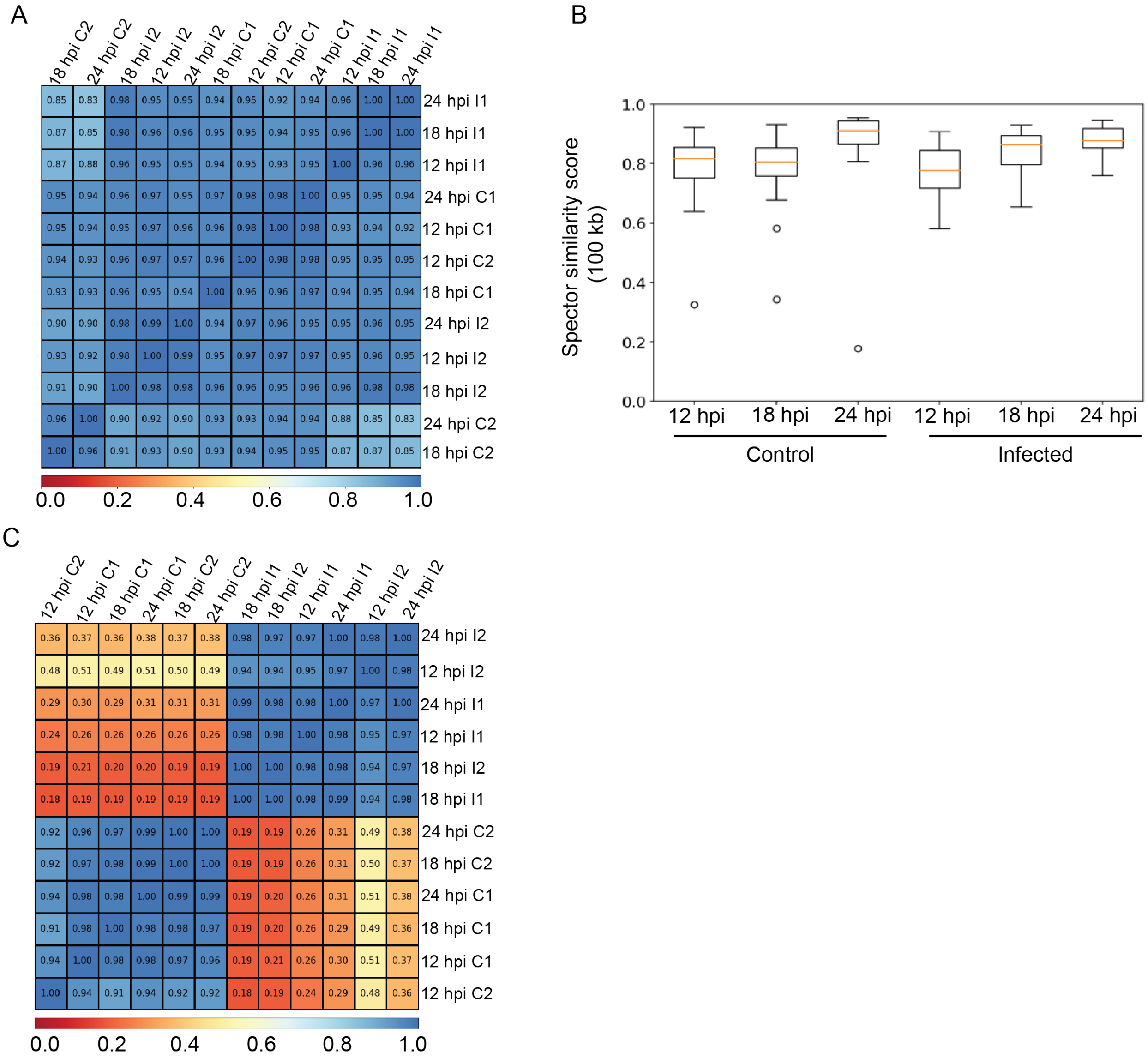


*Figure S 1: A) Genome wide Pearson correlation between ATAC-seq samples at 10 kb resolution. B) Spector similarity score estimated at 100 kb resolution between biological replicates for each condition per time point. C) Genome wide Pearson correlation between RNA-seq samples at 10 kb resolution.*

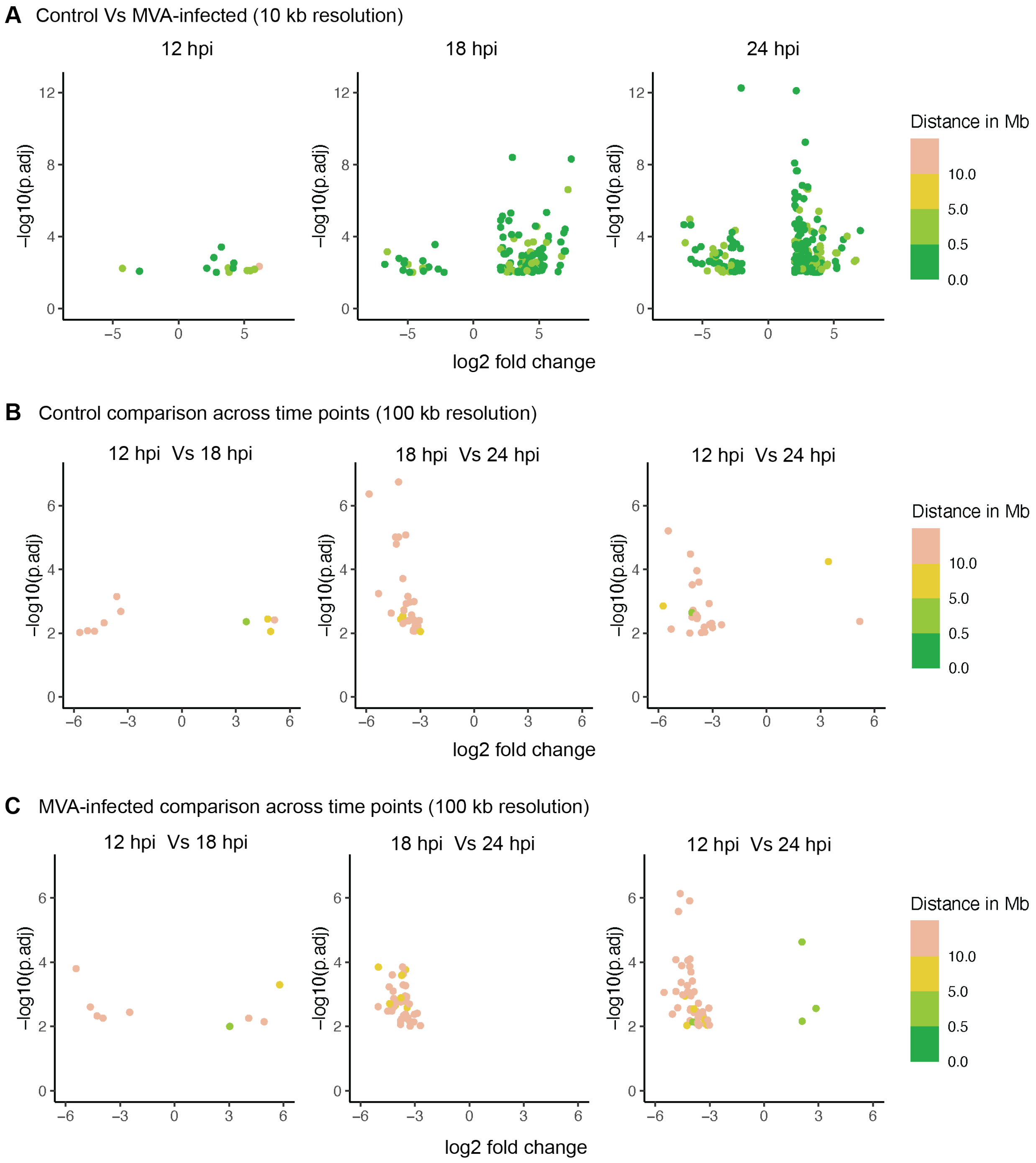


*Figure S 2: A) Genomic regions with significant differences in the number of contacts between control and MVA infection were identified using multiHiCcompare at 10 kb resolution and plotted as log2 fold change in contact frequency (x-axis) and negative log10 adjusted p value (y-axis). Color scale annotates the distance between contacting regions. Points in the positive axis represent infection-bias regions (more contacts due to MVA infection than in control) and points in the negative axis represent control-bias regions. Genomic regions with significant differences in the number of contacts between time points B) in control cells C) in MVA-infected cells were identified using multiHiCcompare at 100 kb resolution and plotted as log2 fold change in contact frequency (x-axis) and negative log10 adjusted p value (y-axis). Color scale annotates the distance between contacting regions. Points in the positive axis represent regions with more contacts in an advanced time point and vice versa.*

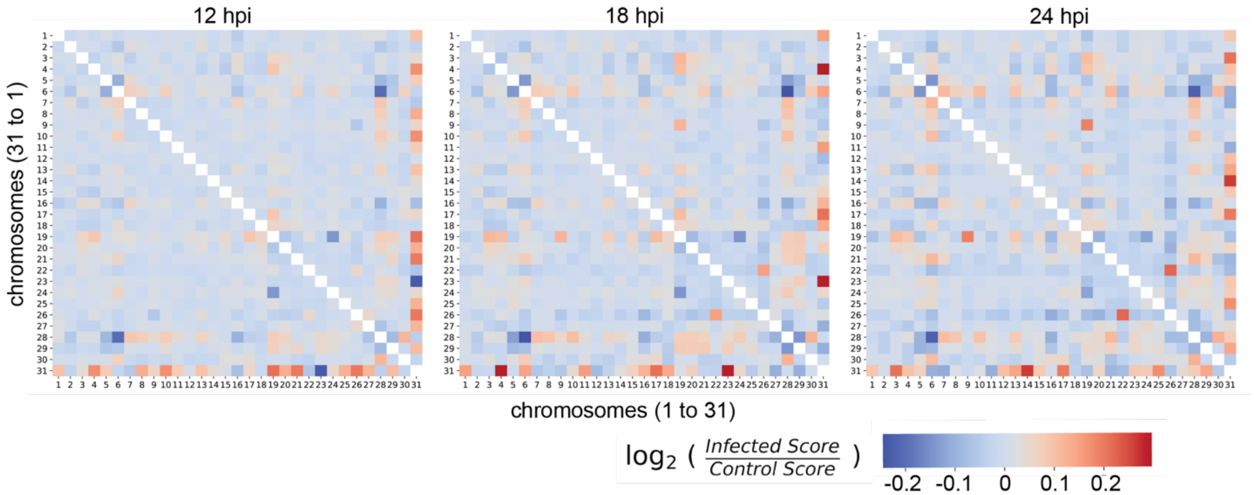


*Figure S 3: Heat map representing differential contacts between pairs of chromosomes at all three timepoints. Color scale represents log2[infected score/control score]. Chromosomes pairs that have increased contacts in infected cells are colored in shades of red and pairs that have increased contacts in control cells are colored in shades of blue.*

| Replicate | Sample | Loops identified with resolution | | | Total |
|---|---|---|---|---|---|
| | | 5 kb | 10 kb | 25 kb | |
| 1 | 12 hr Control | 2888 | 3749 | 3509 | 10146 |
| 2 | 12 hr Control | 8653 | 5615 | 3399 | 17667 |
| 1 | 12 hr Infected | 2007 | 3183 | 3581 | 8771 |
| 2 | 12 hr Infected | 7173 | 5119 | 3683 | 15975 |
| 1 | 18 hr Control | 4897 | 4726 | 3946 | 13569 |
| 2 | 18 hr Control | 8127 | 5444 | 3845 | 17416 |
| 1 | 18 hr Infected | 2902 | 3497 | 3665 | 10064 |
| 2 | 18 hr Infected | 6705 | 4702 | 3972 | 15379 |
| 1 | 24 hr Control | 8672 | 5603 | 3596 | 17871 |
| 2 | 24 hr Control | 7954 | 5376 | 3762 | 17092 |
| 1 | 24 hr Infected | 6229 | 4803 | 3686 | 14718 |
| 2 | 24 hr Infected | 5679 | 4419 | 3890 | 13988 |

*Table S 4: Loop summary*

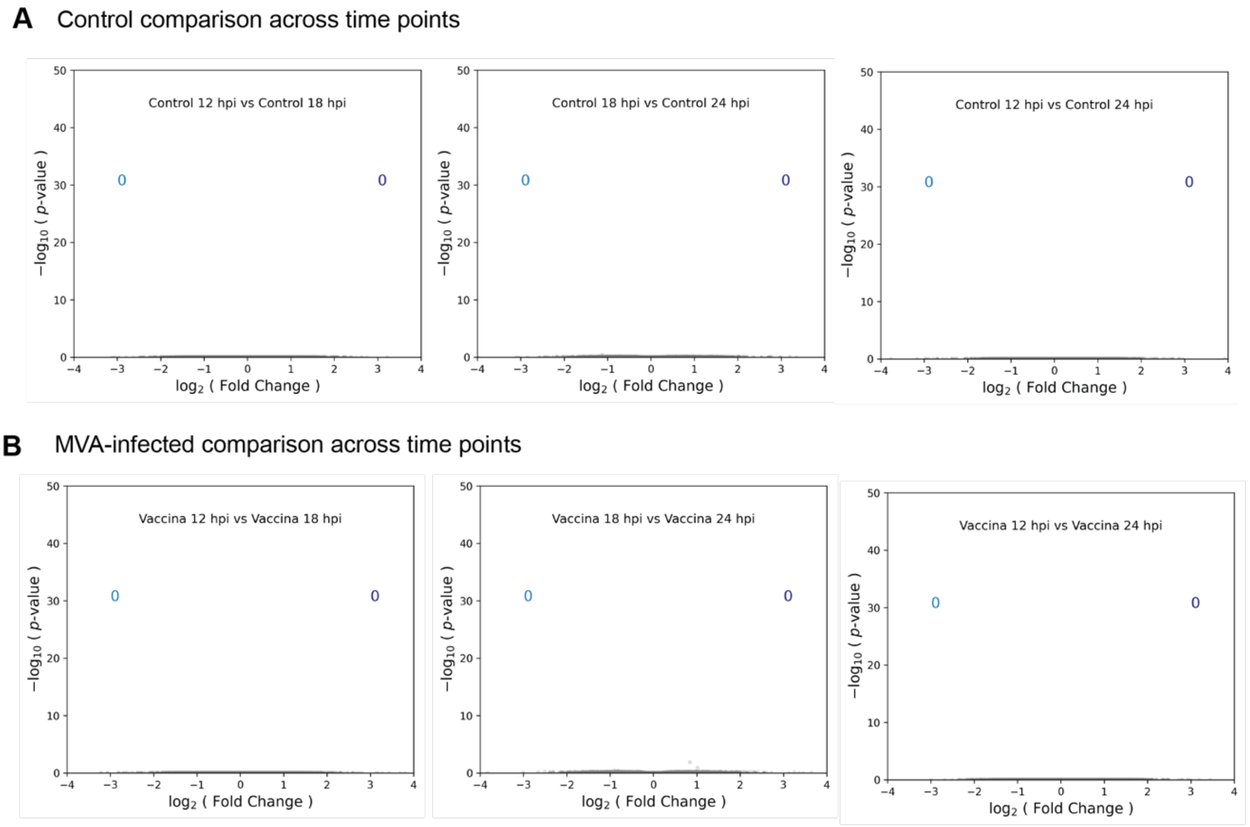


*Figure S 4: Volcano plots represent loop comparison between time points in A) control and B) MVA-infected cells. Grey dots represent all called loops however, zero differential (adjusted p value <0.05) loops were identified in all pair wise comparisons.*

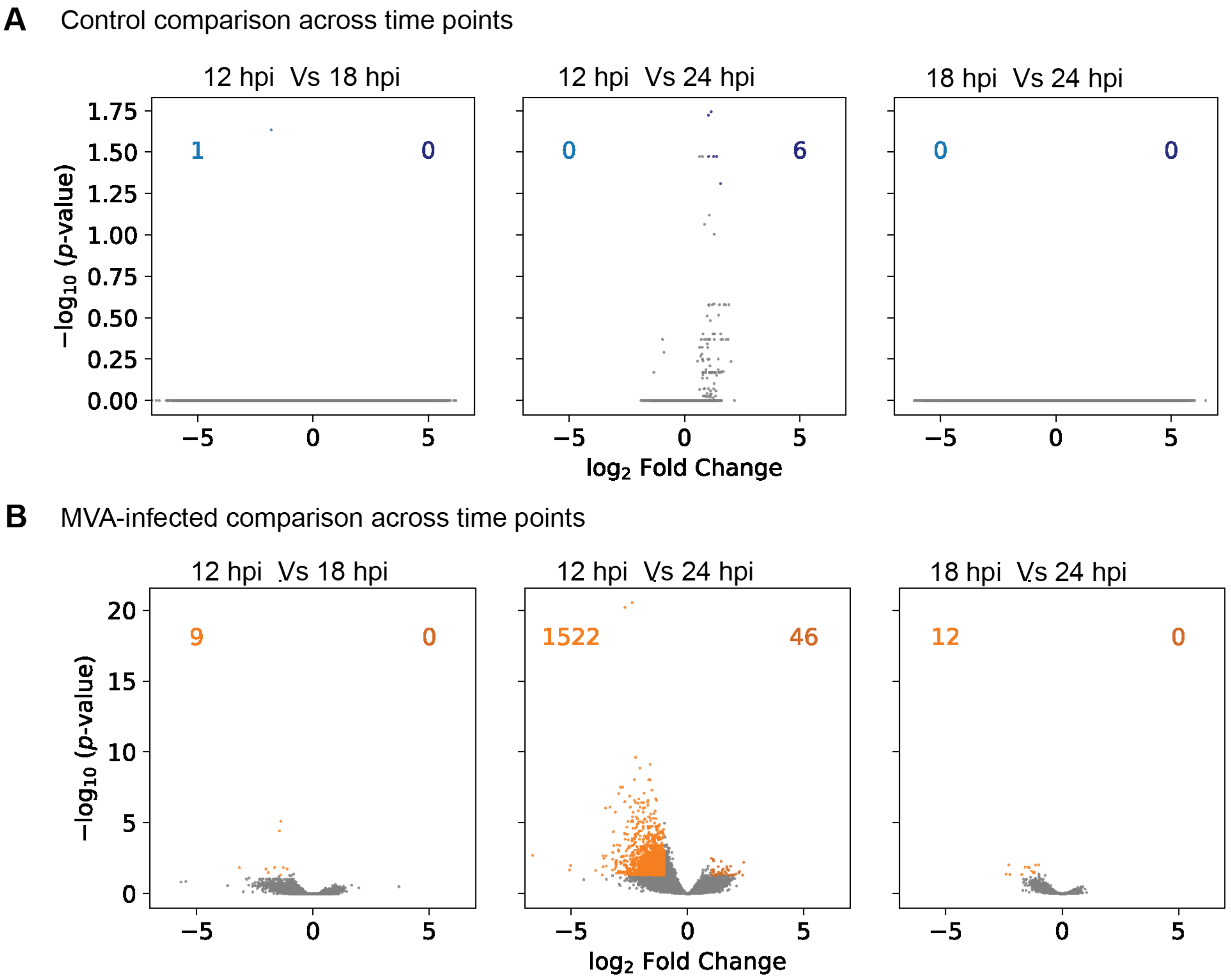


*Figure S 5: Volcano plots demonstrate differentially accessible OCRs between time points in A) mock control and B) MVA-infected Vero cells. Grey dots represent all OCRs. Significantly different (adjusted p value<0.05) OCRs with log2 fold change< -1 or >1 are colored. Points in the positive axis represent differential OCRs with accessibility in an advanced time point and vice versa.*

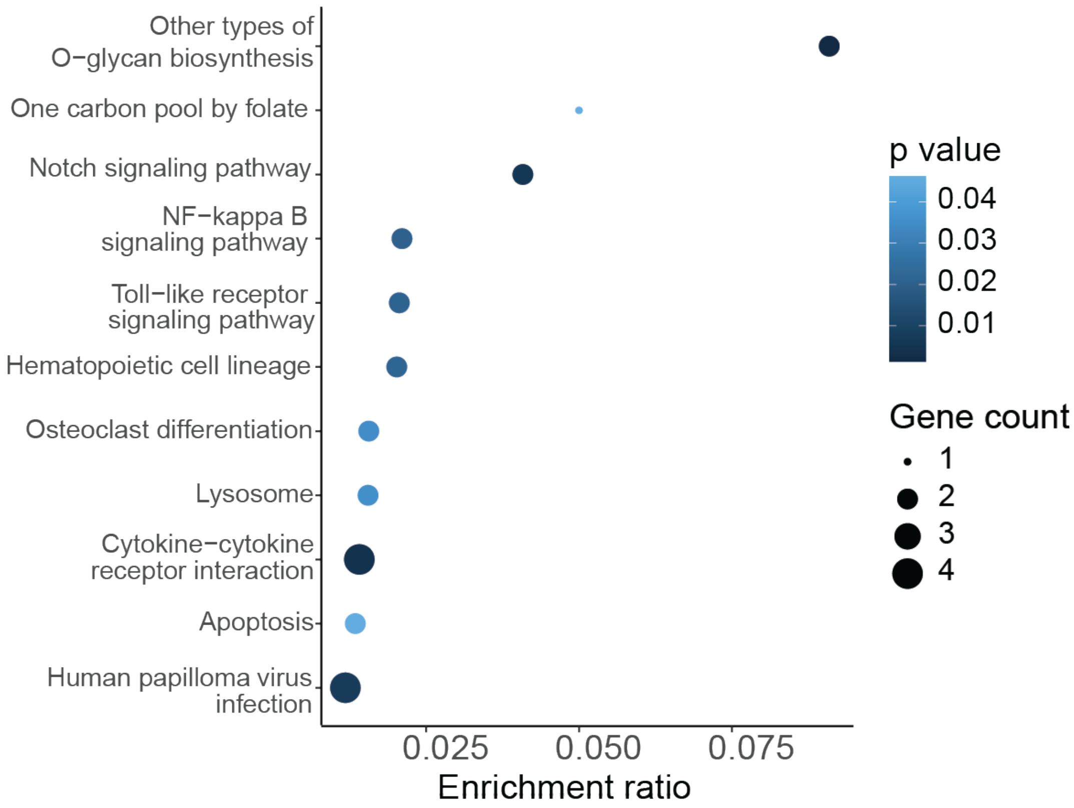


*Figure S 6: Gene regulatory pathways that are potentially suppressed due to MVA infection, as predicted by Kegg Orthology analysis of control-biased differentially accessible genes. Pathways are shown in the decreasing order of their gene enrichment ratio.*

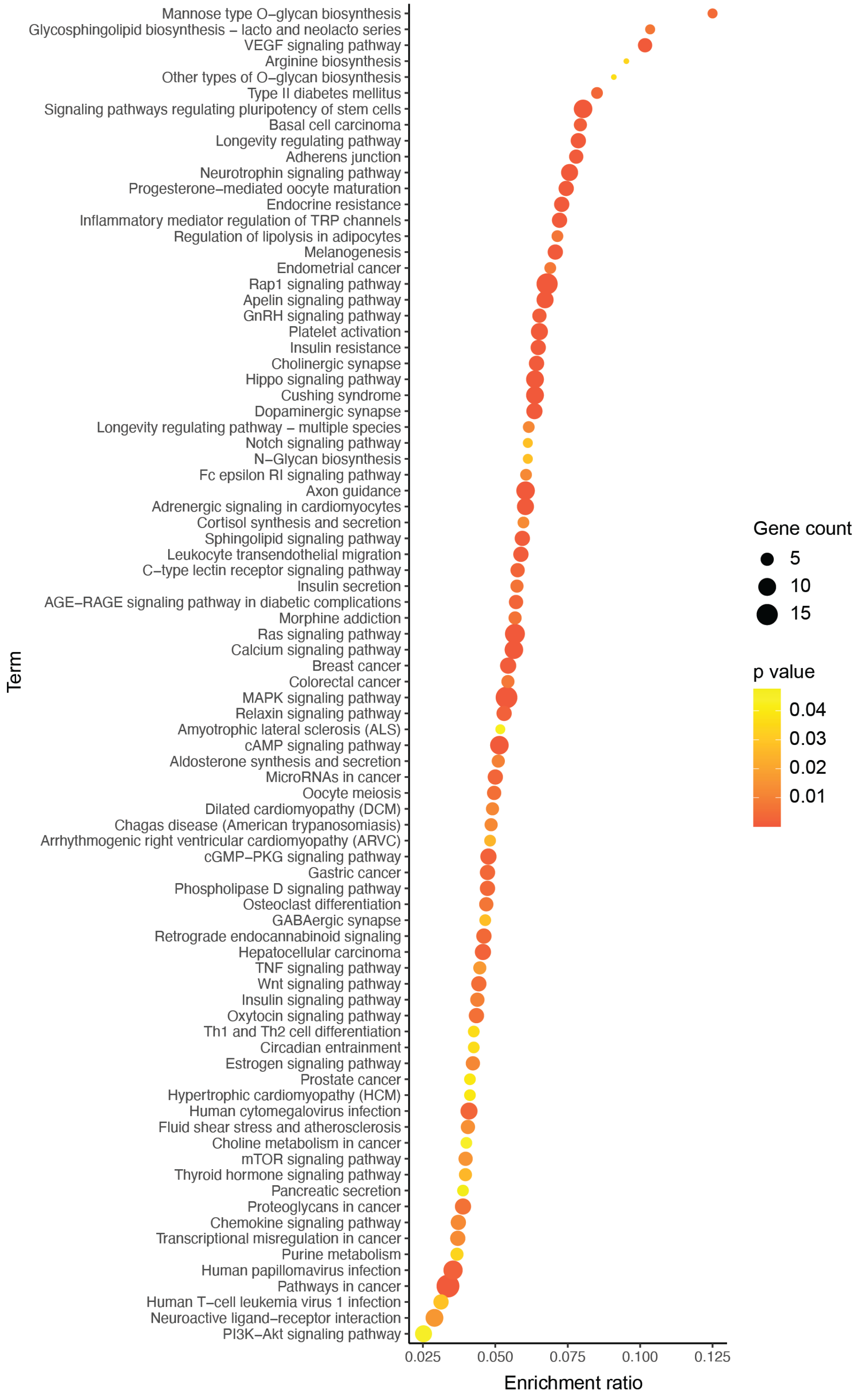


*Figure S 7: Gene regulatory pathways that are potentially activated due to MVA infection, as predicted by Kegg Orthology analysis of infection-biased differentially accessible genes. Pathways are shown in decreasing order of their gene enrichment ratio.*

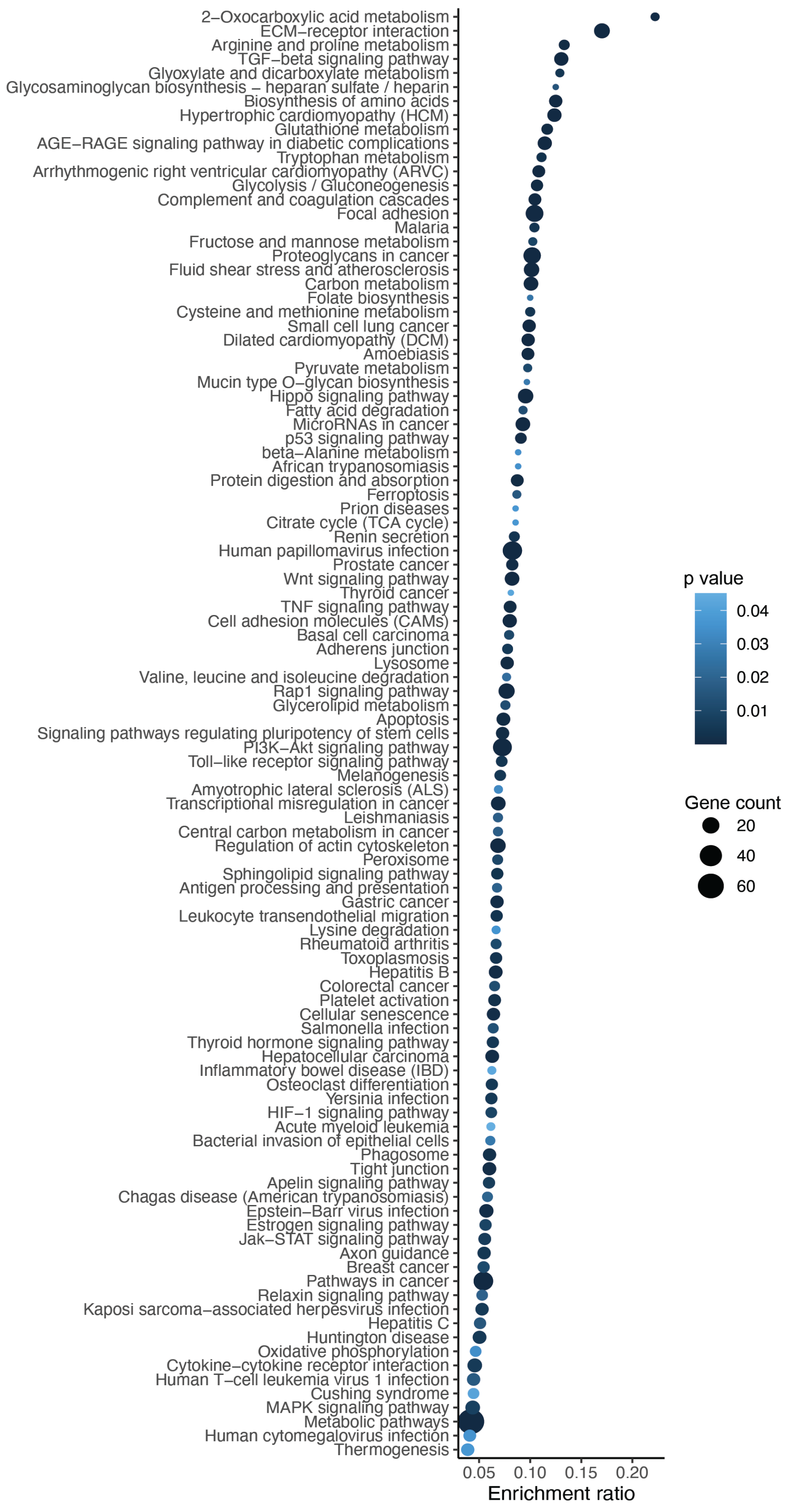


*Figure S 8: Gene regulatory pathways that are potentially suppressed due MVA infection, as identified by Kegg Orthology analysis of downregulated genes. Pathways are shown in the decreasing order of their gene enrichment ratio. (Extended list of main text Figure 6A, left panel).*

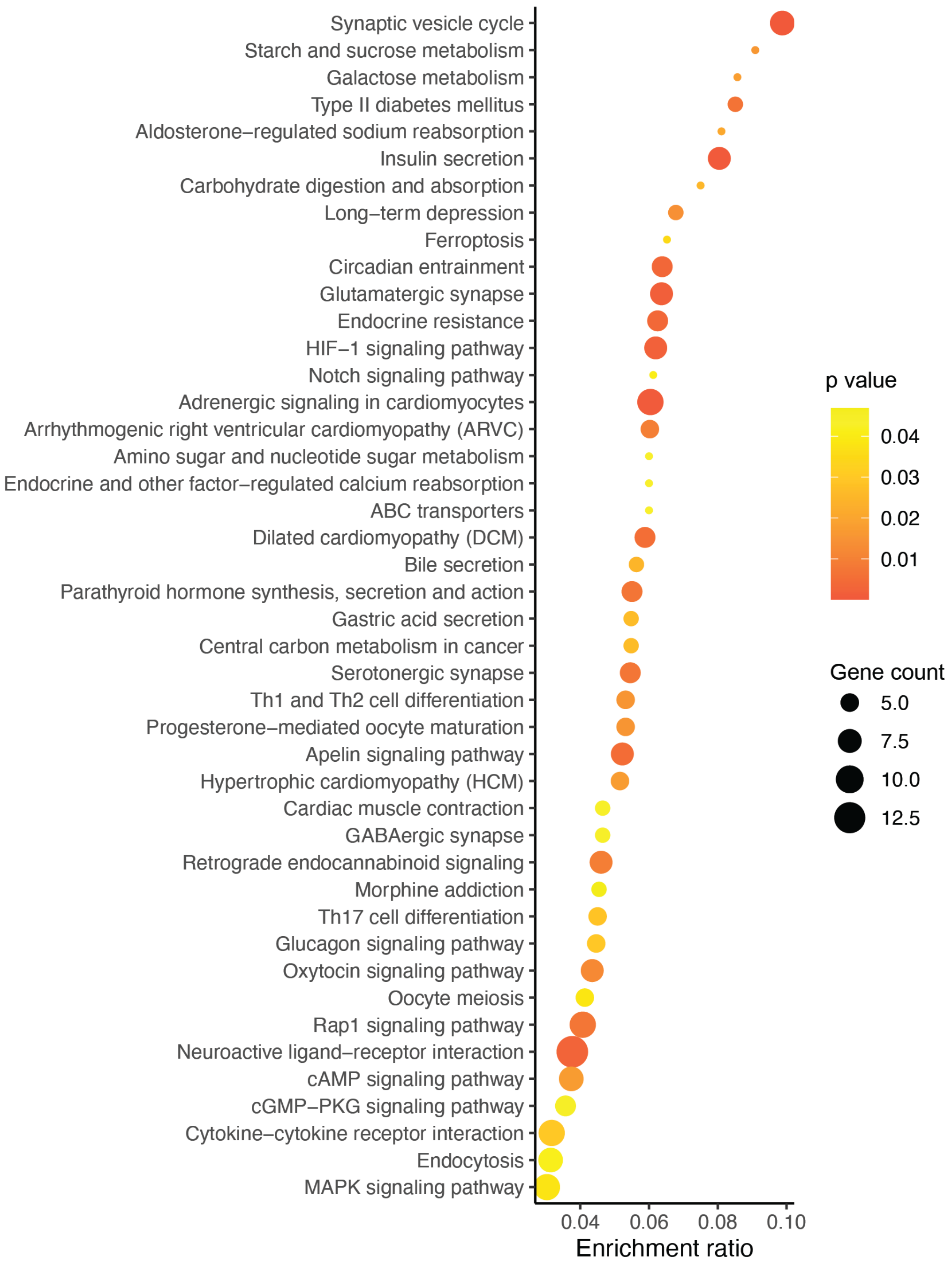


*Figure S 9: Gene regulatory pathways that are potentially activated due to MVA infection, as identified by Kegg Orthology analysis of upregulated genes. Pathways are shown in the decreasing order of their gene enrichment ratio. (Extended list of main text Figure 6A, right panel)*
